# Consumption of artificially sweetened beverages during pregnancy impacts infant gut microbiota and body mass index

**DOI:** 10.1101/2020.04.20.050195

**Authors:** Isabelle Laforest-Lapointe, Allan B. Becker, Piushkumar J. Mandhane, Stuart E. Turvey, Theo J. Moraes, Malcolm R. Sears, Padmaja Subbarao, Laura K. Sycuro, Meghan B. Azad, Marie-Claire Arrieta

## Abstract

Artificial sweetener consumption by pregnant women has been associated with an increased risk of infant obesity, but the underlying mechanisms are unknown. We aimed to determine if maternal consumption of artificially sweetened beverages (ASB) during pregnancy is associated with modifications of infant gut bacterial community composition during the first year of life, and whether these alterations are linked with infant body mass index (BMI) at one year of age. This research included 100 infants from the prospective Canadian CHILD Cohort Study, selected based on maternal ASB consumption during pregnancy (50 non-consumers and 50 daily consumers). We identified four microbiome clusters, of which two recapitulated the maturation trajectory of the infant gut bacterial communities from immature to mature and two deviated from this trajectory. Maternal ASB consumption was associated with the depletion of several Bacteroides sp. and higher infant BMI. As we face an unprecedented rise in childhood obesity, future studies should evaluate the causal role of gut microbiota in the association between maternal ASB consumption, infant development and metabolism, and body composition.

## INTRODUCTION

Childhood obesity in the United States increased from 5 to 18.5 percent between 1978 and 2016^1^, magnifying the risk of cardiometabolic disease and mental health disorders later in life^2^. Recent work from the CHILD Cohort Study showed that maternal consumption of artificially sweetened beverages (ASB) during pregnancy is associated with higher infant body mass index (BMI) at one year of age^3^. Importantly, this association was independent of key obesity risk factors, such as maternal BMI, smoking, poor diet, diabetes, short breastfeeding duration, and earlier introduction of solid food^3^. Similar associations have been reported in several other prospective birth cohorts^4^, but the underlying mechanism has not been studied.

The gastrointestinal tract, a key site for host metabolic regulation^5,6^, is colonized by a vast community of microbes including bacteria, viruses, and micro-eukaryotes^7^. The gut microbiome is highly heterogeneous during infancy, characterized by colonization patterns^8–10^ that are influenced by the maternal microbiome^11,12^, method of birth ^13–15^, infant nutrition (breast milk or formula)^16–18^, and antibiotic treatment^14,19^. Simultaneously, important aspects of metabolic development occur during this period of life, many of which rely on interactions between microbes and host cells^20^. Recent studies in mice show that artificial sweetener consumption during pregnancy predisposes offspring to increased weight gain through behavioral (i.e. preference for sweet foods, appetite increase) and physiological mechanisms (i.e. stimulation of intestinal sugar absorption, increased postnatal weight gain, altered lipid profiles, downregulation of hepatic detoxification, and increased insulin resistance)^21–24^. Suez *et al.*^25^ demonstrated that artificial sweetener consumption in adult mice directly impacts gut microbiome composition and function, leading to an increase in host glucose intolerance. More recently, Stichelen *et al.*^24^ addressed gestational exposure to artificial sweeteners, finding changes in bacterial metabolites and an decrease in *Akkermansia municiphila* in the pups’ gut microbiome. However, the consequences of maternal artificial sweetener consumption during pregnancy on the infant gut microbiota has not been reported in humans.

To address this knowledge gap and build on our prior observations in the CHILD Cohort Study, we evaluated the association of maternal artificially sweetened beverage (ASB) consumption during pregnancy with the infant gut microbiota in a subset of 100 infants (50 with daily maternal ASB consumption during pregnancy and 50 unexposed controls). We employed next generation sequencing of the 16S rRNA amplicon gene combined with a community typing analysis (Dirichlet Multinomial Mixtures [DMM] modelling)^26^ to understand if ASB intake was associated with a shift in infant microbiota composition that might explain the relationship between maternal ASB intake during pregnancy and infant BMI at one year of age.

## METHODS

### Study design and population

We used data and samples collected through the CHILD Cohort Study^27,28^, a Canadian general population birth cohort (3621 families recruited across four provinces) including singleton pregnancies (>35 weeks gestational age with no congenital abnormalities) enrolled from 2008 to 2012. From this cohort, we completed a case-control study by selecting 100 infants divided equally between mothers that reported little or no ASB consumption (less than one per month) or high ASB consumption (one or more per day) during pregnancy. The groups were balanced for six potential confounding factors known to influence the gut microbiome: infant sex, birth mode, breastfeeding at three and 12 months, maternal BMI, and antibiotic use in infants before 12 months (antibiotics before three months old was an exclusion criterion; eTable 1). To characterize the gut microbiome, stool samples were acquired at three and 12 months of age for a total of 200 samples. This study was approved by the University of Calgary Conjoint Health Research Ethics Board (CHREB) and ethics committees at the Hospital for Sick Children, and the Universities of Manitoba, Alberta, and British Columbia. Written informed consent was obtained from mothers during enrollment to the CHILD Study.

### Maternal diet in pregnancy

Maternal dietary assessment in pregnancy has previously been described^3^. Briefly, a food frequency questionnaire (FFQ) was completed during the second or third trimester and ASB consumption was evaluated using reports of “diet soft drinks or pop” (i.e. soda) (serving = 12 oz / one can) and “artificial sweetener added to tea or coffee” (serving = 1 packet). Other dietary variables included: sugar-sweetened beverages, Healthy Eating Index (HEI) total score (see eMethods), added sugar and total energy intake.

### Infant BMI

BMI was measured by CHILD staff to the nearest 0.1 kg around one year of age (mean = 12.0 months ± 0.8 [sd]) and height to the nearest 0.1 cm. Age- and sex-specific BMI-for-age z-scores were calculated following the World Health Organization reference^29^.

### Other variables

The following variables were considered in univariable analyses (see eMethods): (1) infant’s sex, age at sample collection, breastfeeding duration (BF duration; months), breastfeeding status at three months (BF at 3M; yes or no), diet at three and six months (Diet at 3M and Diet at 6M; both defined in 8 categories allocated based on the presence in the infant’s diet of breastfeeding, formula, and solids), solids at three and six months (Solids at 3M and Solids at 6M), formula feeding at three months (FF at 3M), number of antibiotic treatments received from six to twelve months (Child 6-12 abx), and secretor status (determined from the single nucleotide polymorphism rs601338 in the *FUT2* gene); (2) mother’s gestational diabetes, age, ethnicity, education, oral antibiotics received pre-delivery (Mother pre-delivery abx), intrapartum antibiotics (Mother intrapartum abx), and secretor status (rs601338 SNP); (3) study site, presence of cats, dogs, and older siblings in the house.

### Stool samples DNA extraction and sequencing

We extracted gut microbial DNA from fecal samples using the DNeasy PowerSoil kit (QIAGEN) according to the manufacturer’s instructions. and amplified the V4 region of the 16S rRNA gene to generate ready-to-pool dual-indexed amplicon libraries as described previously^30^ (see eMethods). Using the DADA2^31^ pipeline, the final dataset contained 4,553,000 quality sequences, a mean (range) of 6,509 (22,995 - 68,265) sequences per sample identified as 954 unique bacterial Amplicon Sequence Variants (ASVs). Samples contained a mean of 40 (10-95) unique ASVs per samples.

### Statistical analysis

We used Dirichlet Multinomial Mixtures (DMM) modelling^26^ on 16S rRNA gene sequencing data to identify clusters of similar bacterial community structure amongst our samples (a technique known as community typing analysis, increasingly used in human microbiome studies^10,32–34^). This technique is increasing employed in microbiome studies for three reasons: (1) identification of unique microbial clusters is unsupervised; (2) cluster size depends on metacommunity variability; and (3) adequate explicit probabilistic model penalises model complexity to optimize cluster number. The lowest Laplace approximation grouped our samples in four unique clusters (Figure 1-2 and eFigure 1).

**Figure 1.**
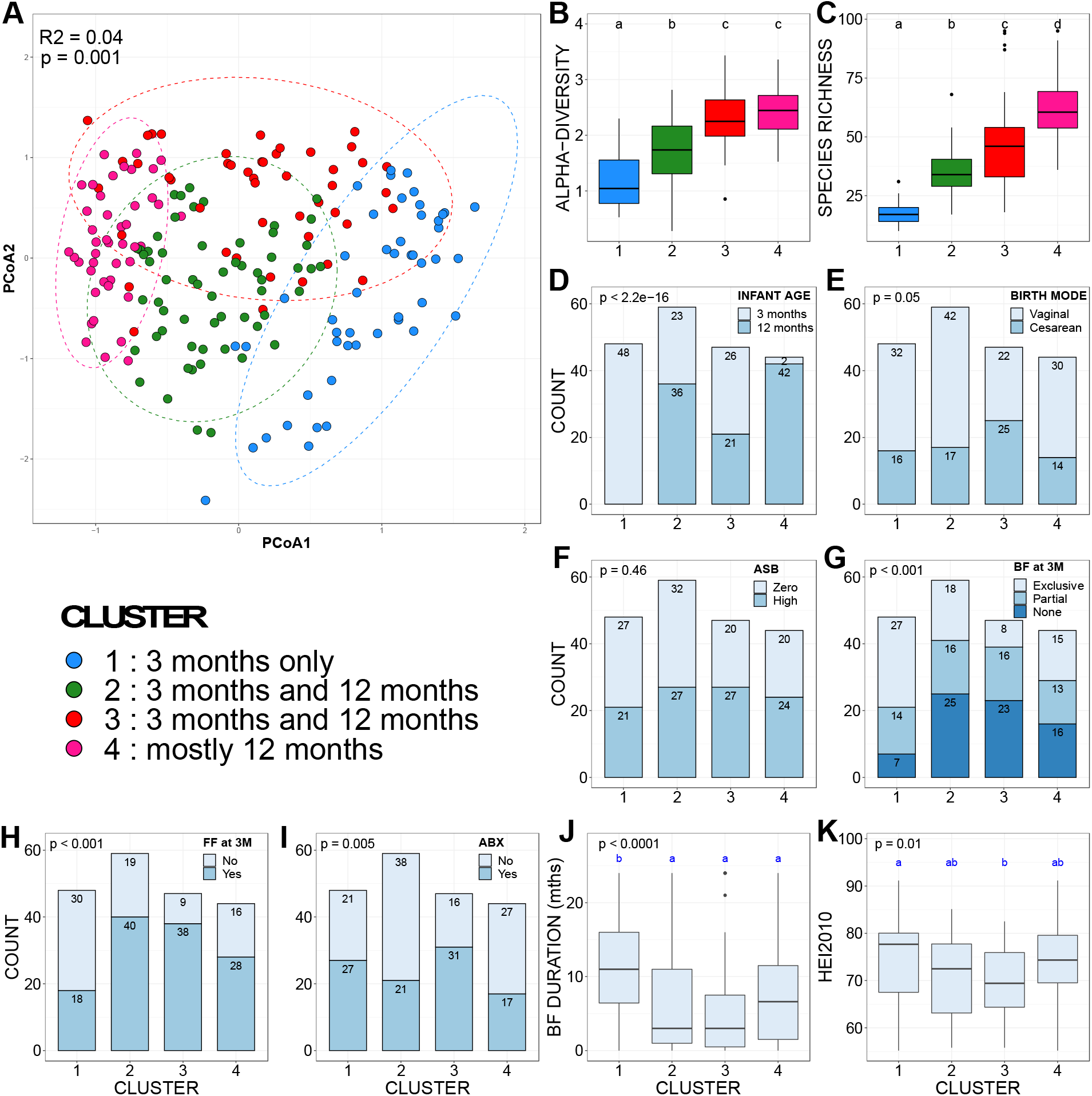
Discrepancies in covariate distribution, alpha- and beta-diversity between clusters. (A) Principal component analysis (PCoA) ordinations of variation in beta-diversity of infant gut bacterial communities based on Bray-Curtis dissimilarities among samples. Ellipses represent 95% confidence intervals. (B-C) Box plots showing the alpha-diversity (richness and Shannon’s diversity) per DMM cluster. The central line denotes the median, the boxes cover the 25th and 75th percentiles, and the whiskers extend to the most extreme data point, which is no more than 1.5 times the length of the box away from the box. Points outside the whiskers represent outlier samples. Letters denoted significant differences (non-parametric Kruskal-Wallis test followed by post-hoc test of Dunn with FDR correction following Benjamini-Hochberg method; P<0.05). (D-K) Variable distribution between clusters tested with non-parametric Kruskal-Wallis test followed by either a post-hoc generalized linear model (glm) with a binomial/logistic distribution (D-I) or (J-K) a post-hoc Dunn test with FDR correction following Benjamini-Hochberg method. Minuscule letters indicate statistical differences between clusters from post-hoc generalized linear model (glm) with a binomial/logistic distribution. “BF at 3M” stands for “breastfeeding at three months” and “FF at 3M” for “formula feeding at three months”. Aside from maternal ASB consumption (F), only the variables that showed a statistical difference in distribution between clusters are presented. No differences were found for maternal age, ethnicity, education, diabetes; study site, household pets, siblings, or introduction of solid foods at 3 or 6 months. Cluster 1 included 48 samples from 48 infants; cluster 2 included 59 samples from 49 infants; cluster 3 included 47 samples from 39 infants; and cluster 4 included 44 samples from 43 infants. See methods for definition of variables.

**Figure 2.**
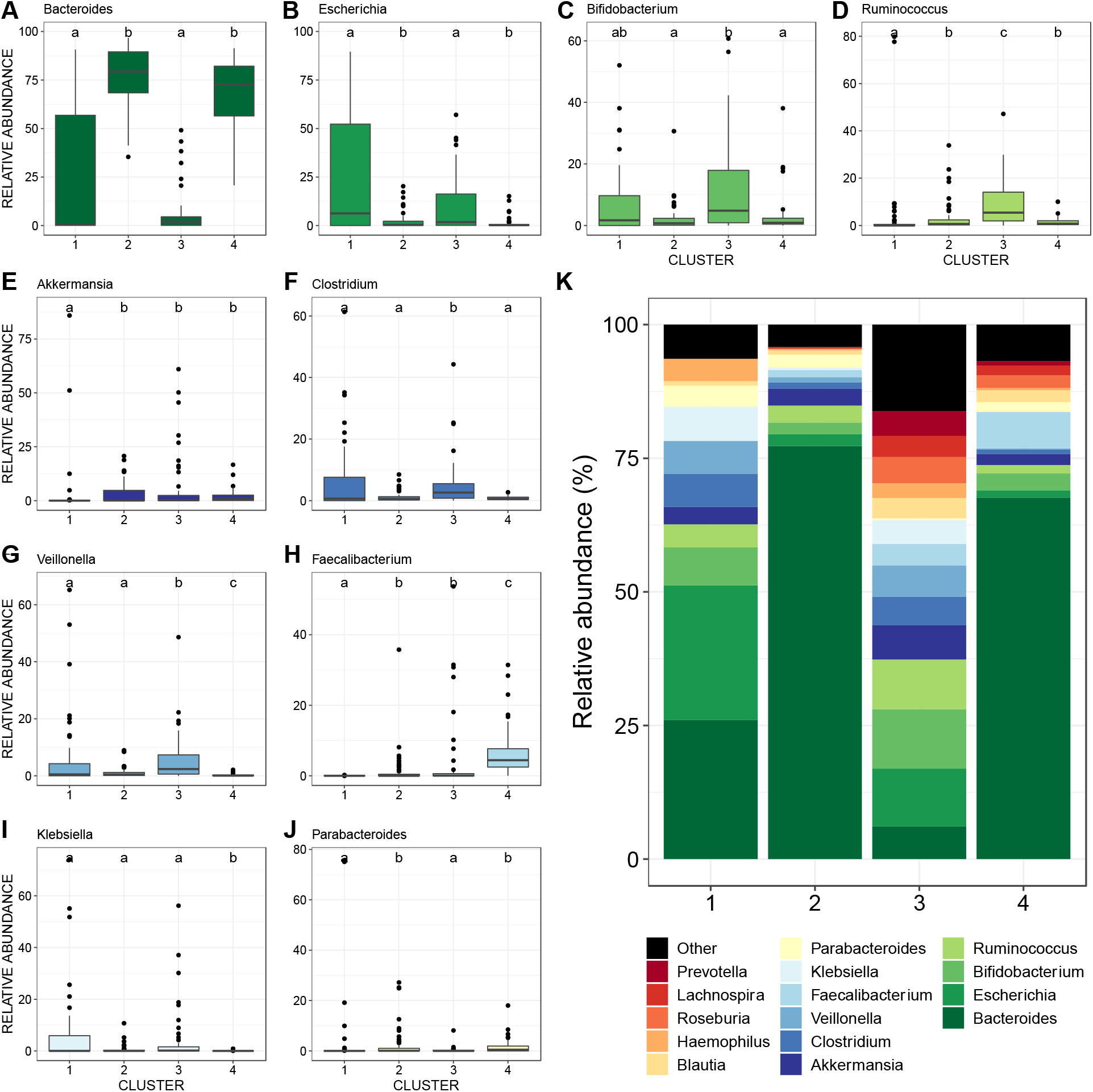
Differences in relative abundances of the dominant bacterial genera between clusters. (A-J) Relative abundance across DMM clusters of the ten most dominant bacterial genera and (K) of the 15 most dominant bacterial genera. Letters indicate significant differences between clusters (non-parametric Kruskal-Wallis test, post-hoc Dunn test with Benjamini-Hochberg FDR correction). Cluster 1 contains only three months of age. Cluster 2 and 3 are composed of a mix three and twelve months of age, and Cluster 4 only 12M (except two samples).

The distribution of variables as well as the variation in bacterial richness (Chao 1), alpha-diversity (Shannon index), and community evenness (Shannon index / log_n_(species richness)) across the DMM clusters were examined by non-parametric Kruskal-Wallis tests followed by post-hoc Dunn tests or generalized linear models (glm) with a binomial/logistic distribution. To explore the changes in taxonomical community structure at a fine scale, we tested for significant differences in the relative abundance of the 10 most dominant bacterial genera across clusters using non-parametric Kruskal-Wallis tests followed by post-hoc Dunn tests with Benjamin-Holmes False Discovery Rate (FDR) correction. To account for potential heteroskedasticity in bacterial community dispersion between groups and avoid the loss of information through rarefaction^35^, we performed a variance stabilizing transformation^35,36^ prior to any statistical tests on beta-diversity. To select variables that could be drivers of infant gut bacterial community structure, we tested for correlations between our variables and community scores on the Principal Component Analysis (PCoA) ordination axes in univariable models (*envfit* function of vegan^37^). The relative influence of the significant drivers of gut bacterial community structure was then assessed statistically in multivariate models using a Permutational Multivariate Analysis Of Variance (PERMANOVA; *adonis* function of vegan^37^) with 999 permutations and visualized using PCoAs based on Bray-Curtis dissimilarities. We used DESeq2 to test for differentially abundant bacterial taxa according to maternal ASB consumption on the 100 most relatively abundant bacterial taxa to limit spurious significance driven by very rare ASVs. Finally, we used linear models on the three- and twelve-months-old samples to test for the influence of maternal ASB consumption and microbial ordination axes (PCoA1 and PCoA2) on infant BMI z-score. The full model’s formula was the following:

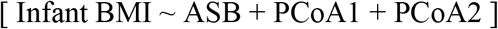

All analyses and graphs were computed in R version 3.6.1 (R Development Core Team; http://www.R-project.org).

## RESULTS

### Microbiome clusters

We performed community typing analysis based on Dirichlet Multinomial Mixtures (DMM) modelling^26^ to identify clusters of similar bacterial community structure amongst our samples. Based on their microbiota composition, the infant fecal samples clustered in four groups (Figure 1-2 and eFigure 1). Gut bacterial species richness (Figure 1B), alpha- (Figure 1C) and beta-diversity (Figure 1A) and taxonomic composition (Figure 2) differed between clusters, reflecting broad community differences. Clusters 1 and 4 comprised microbial communities reflecting the well-described effect of temporal maturation during the first year of life; with cluster 1 comprising only three-month (3M) samples and cluster 4 comprising almost exclusively twelve-month (12M) samples. Clusters 2 and 3 comprised a mixture of 3M and 12M samples. Compared to the other three clusters, cluster 1 showed a higher proportion of exclusive breastfeeding. Cluster 3 included a higher proportion of mothers receiving antibiotics, infants born by C-section and formula feeding (Figure 1). However, there was no difference in maternal ASB consumption between clusters, suggesting that this exposure did not influence the compositional differences that drove cluster classification (Figure 1F). In addition, the clusters did not differ in terms of maternal sugar intake, gestational diabetes, age, parity, ethnicity, education, antibiotics, study site, infant antibiotics, or infant or mother secretor status.

### Relative influence of ASB on microbial community structure

*Envfit* analysis (univariable models) identified thirteen variables as significant drivers of gut bacterial beta-diversity from which we selected eight non-redundant variables to build our models: infant age, maternal intrapartum antibiotics, maternal ethnicity, birth mode, breastfeeding status at three months, presence of older siblings, infant secretor status, and maternal ASB consumption (Figure 3A and eFigure 2). Considering the complete dataset, the significant predictors were infant age, maternal ethnicity, intrapartum antibiotics, and birth mode. The same four variables, plus breastfeeding status at 3 months, were tested in a PERMANOVA (multivariable model), altogether explaining 14.2% of community variance (Table 1). Maternal ASB consumption was a significant predictor of infant gut bacterial composition only in the multivariable model (R^2^ = 0.7%; Table 1). Birth mode (vaginal vs. C-section) had also a significant influence on community composition (R^2^=0.8%), but to a lesser extent than infant age (R^2^ = 7.3%) and mother’s ethnicity (R^2^ = 2.5%; Table 1).

**Figure 3.**
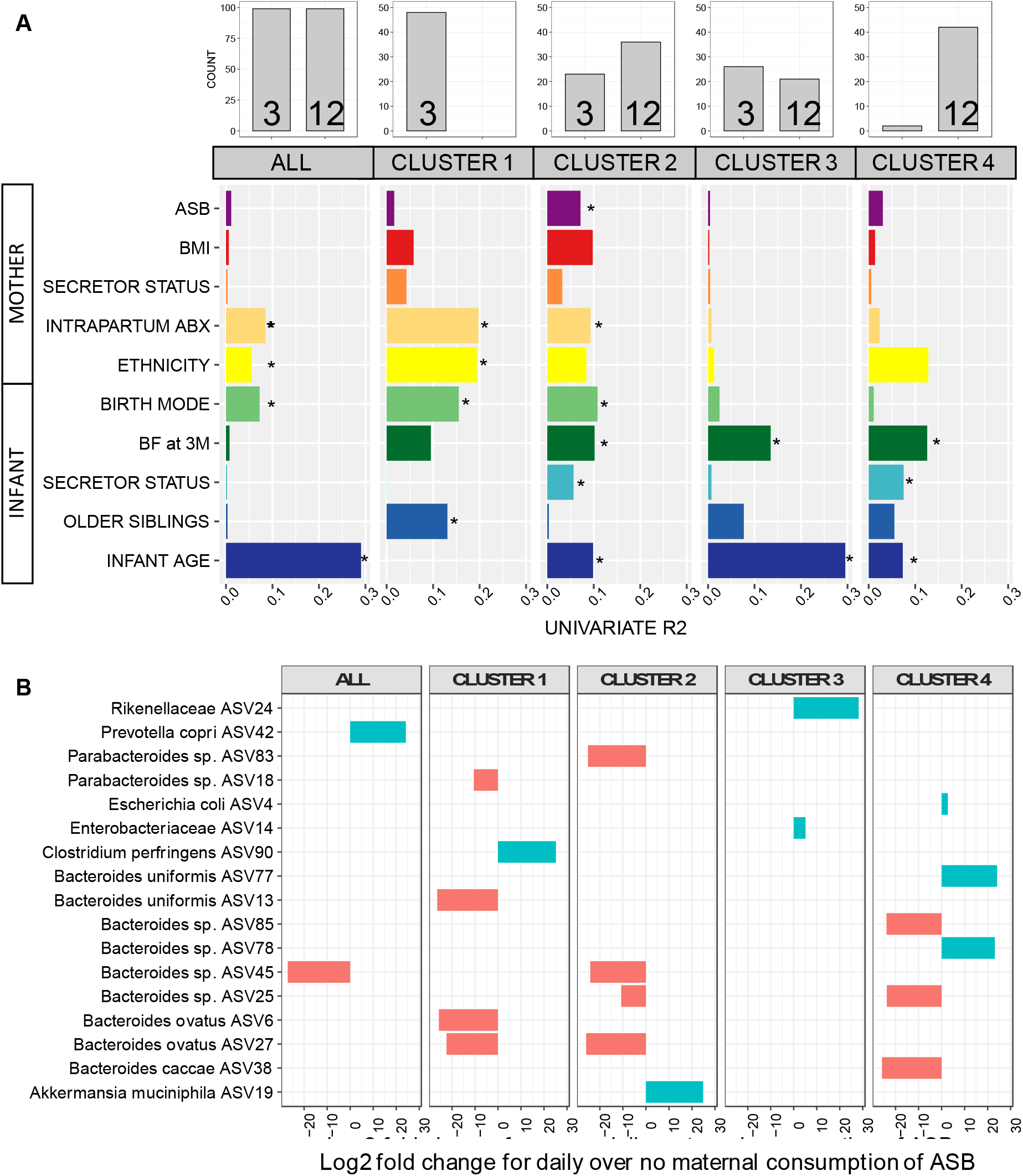
Drivers of gut bacterial beta-diversity and indicator taxa associated with maternal consumption of ASB differ between clusters. (A) Univariate models showing significance and explained variance of 10 variables on bacterial community structure across all data and each cluster subset. Horizontal bars show the amount of variance (R^2^) explained by each covariate in the model as determined by *envfit*. Asterisk denotes the significant covariates in each data subset (P<0.05). All 32 variables considered in this study are shown in eFigure 2. In this figure, ASB represents artificially sweetened beverages and BF at 3M represents infant’s breastfeeding status at three months (see methodology). (B) 14 bacterial taxa identified as significant features associated with maternal consumption of ASB by DESeq2.

**Table 1.**
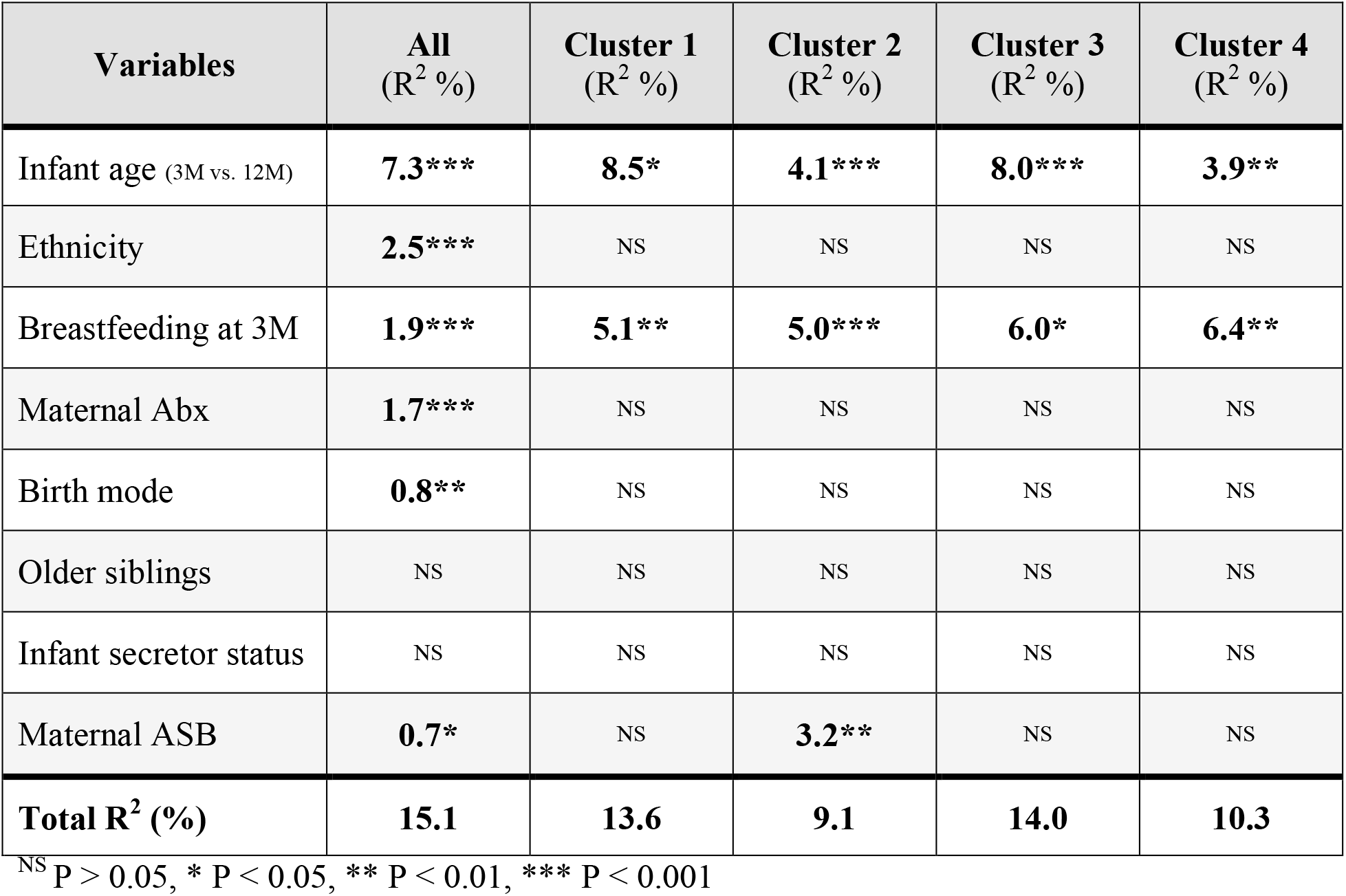
Maternal consumption of ASB during pregnancy is associated with bacterial community assembly during the first year of life. Permutational Analysis of Variance (PERMANOVA) of gut bacterial community composition (Bray-Curtis dissimilarities) testing associations with different explanatory variables (a: all data, b-e: clusters 1-4). The model on the complete dataset (ALL) accounts for repeated measures. The set of variables to be tested was chosen based on results from univariate *envfit* models: infant age, antibiotics received by mother at birth, mother’s ethnicity, birth mode, breastfeeding status at three months, presence of older siblings, and maternal ASB consumption.

Next, we repeated the beta-diversity analyses separately within each of the 4 clusters. *Envfit* univariable models identified distinct drivers for each cluster (Figure 3A). Interestingly, the drivers of beta-diversity in cluster 1 (only 3M samples) were mainly maternal factors (i.e. birth mode, mother’s ethnicity, intrapartum antibiotics) whereas the drivers of cluster 4 (mostly 12M) were infant factors (infant’s secretor status, breastfeeding at three months, and infant age (Figure 3A). Cluster 2 was the only cluster in which maternal ASB consumption was associated with beta-diversity (R^2^ = 3.2%), and this association was confirmed by the univariable (Figure 3A, eFigure 2) and multivariable (Table 1) analyses.

We tested for associations of specific bacterial features in the infant gut with maternal ASB consumption. In the complete dataset, we identified two ASVs associated with maternal consumption of ASB, one species being depleted (*Bacteroides* sp. ASV45, log2 fold change = −27.2 and another species enriched (*Prevotella copri* ASV42, 24.2) among infants exposed to high maternal ASB intake (Figure 3B). Repeating this test within each cluster, we identified 15 additional ASVs enriched or depleted. For cluster 2, one ASV was enriched (ASV19, *Akkermansia municiphila*, 24.9) and four depleted (*Bacteroides ovatus* ASV27, −25.9; Parabacteroides sp. ASV83, −25.2; *Bacteroides* sp. ASV45, −24.9; *Bacteroides* sp. ASV25, −10.7) with maternal ASB consumption (Figure 3B). All adjusted p-values were below 0.001.

### Association of ASB and the microbiome with infant BMI at one-year-old

Finally, using a multivariable linear model on the complete dataset, we tested the association of maternal ASB consumption and microbial community composition with infant BMI z-score at one year of age. Our results confirmed that daily maternal ASB consumption is associated with higher infant BMI (ß-estimate = 0.42, 95%CI 0.03:0.80, P = 0.037; Table 2), and showed that BMI was associated with the microbiome composition at 12 months (PCoA1 axis; ß-estimate = − 0.71, 95%CI −1.40:-0.01, P = 0.048; Table 2) but not at three months (not shown). These results suggest that features of PCoA1 (i.e. lower relative abundance of *Bacteroidetes* and *Faecalibacterium*, and higher relative abundance of *Escherichia*, *Klebsiella*, *Bifidobacterium*, *Haemophilus*, *Clostridium*, and *Veillonella*; eFigure 3) are inversely associated with infant BMI.

**Table 2.** Maternal consumption of ASB during pregnancy is associated with higher infant BMI at one-year-old. Linear model showing the explanatory power of maternal ASB consumption on infant BMI z-score at one year old, as well as the two main axes of ordination of bacterial community structure (beta-diversity) on samples acquired at three and twelve-month-old. The full models are:

| Variables | Infant BMI z-score at 1 year |  |  |  |
| --- | --- | --- | --- | --- |
| | β-est. | 95% CI | P-value | R<sup>2</sup> |
| Maternal ASB (daily vs. no consumption) | **0.42** | **[0.03,0.81]** | **0.037** | **4.1%** |
| *12 months microbiome* |  |  |  |  |
| PCoA axis 1 | **-0.71** | **[-1.40, -0.01]** | **0.048** | **3.9%** |
| PCoA axis 2 | NS | NS | NS | NS |
| **Total adj. R<sup>2</sup>** |  | **8.1%** |  |  |

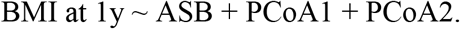

Microbial variables were transformed (squared root and order quantile normalized respectively) to achieve normality. Here we present only the best model for 12 months fitted by stepwise selection by Akaike information criterion because we detected no association between BMI at one year old and microbiota composition at three months old.

## DISCUSSION

In defining links between maternal ASB consumption and infant BMI, we provide new evidence suggesting that maternal consumption of ASB during pregnancy (1) influences the establishment of the infant gut microbiome, particularly in infants diverging from what has previously been described as the typical microbiome maturation trajectory (Table 1, Figure 3A); and (2) is associated with an increase in infant BMI at one-year-old (Table 2). To our knowledge, this is the first human study to report the impact of maternal consumption of ASB on the infant gut microbiome, and its potential influence on infant BMI. In light of recent data showing that ASB can drive dysregulation of energy metabolism in mice through changes in the gut microbiome^24,25,38,39^, our study suggests that infants exposed to ASB through their mothers may be at higher risk of shifts in microbial community structure related to early-life predisposition to metabolic diseases^40,41^.

In our study, broad shifts in bacterial community structure were significantly associated with infant BMI at one-year-old. We also identified 9 bacterial taxa from *Bacteroides* sp. that were enriched (3 ASVs) or depleted (6 ASVs) at high levels of maternal ASB consumption, suggesting a mechanism of influence on infant weight gain involving specific taxa of the gut microbiome. The taxa *Akkermansia municiphila* and genus *Bacteroides* have previously been identified by various studies to be respectively decreased and enriched as a consequence of ASB consumption^25,38,39,42^. Our results differ from previous findings for *A. municiphila* and suggest that *Bacteroides* patterns of enrichment or depletion might be species- or strain-specific, warranting further research with deeper resolution.

As reported by Bian *et al.*^38,39^ in two studies with adult mice, and by Nettleton *et al.*^43^ in a study on dams and their offspring, ASB have been shown to alter gut bacterial community composition (increase of *Bacteroides* and reductions of *Lactobacillus* and *Clostridium*) and increase body weight in parallel with an enrichment of energy metabolism bacterial genes. The functional cluster analyses by Bian *et al.*^38,39^ revealed activation of genes related to carbohydrate absorption and increases in metabolic pathways related to glycolysis and sugar and xylose transport^38^. Sucralose treatment resulted in an increase in bacterial pro-inflammatory mediator genes in mice^39^. Likewise, Chi *et al.*^42^ found that consumption of the artificial sweetener neotame altered the alpha- and beta-diversity of mice gut microbiome, and led to a decrease in butyrate synthetic genes and changes to the fecal short chain fatty acids cluster. Overall, accumulating evidence suggests that the alterations of host gut bacterial community structure through the consumption of ASB is reflected in bacterial and host metabolic gene clusters, which might explain the increase in weight gain. Based on this evidence and our current results, we hypothesize that gestational exposure to ASB impacts infant gut bacterial communities either indirectly through disruption of vertical transmission of the maternal microbiome, or directly through lactation during breastfeeding. However, our study is underpowered to definitively assess whether gut microbiome mediate the relationship between maternal ASB and infant BMI. Additional work including functional evidence from metagenomics and metabolomics will determine if the bacterial taxa and compositional changes associated with high maternal ASB consumption in our study are causally implicated in energy metabolism dysregulation and infant body composition.

Overall, our study validates previous findings^3^ that maternal consumption of artificial sweeteners is associated with a higher BMI at one-year-old, and provides unique and timely evidence that the infant gut microbiome could play a role in this effect, especially for susceptible infants displaying a disrupted maturation trajectory (reduced alpha-diversity and species richness) of their gut microbiome and a high relative abundance of *Bacteroides*. Our study also confirms recent descriptions of infant microbiome development and confirms the influence of several known determinants of the gut microbiome during the first year of life^11–14,16,17,19^ including maternal antibiotics, breastfeeding, birth mode and ethnicity.

The major strength of our study is the combination of state-of-the-art community typing analysis of the gut bacterial communities combined with the standardized prospective evaluation of maternal ASB consumption. Limitations of our study lie in risk of measurement error in self-reported dietary exposures and our inability to distinguish between different types of ASB or account for artificial sweeteners in foods. Also, we did not assess maternal diet after delivery, so we could not directly investigate the impact of prenatal ASB exposure *in utero* versus postnatal exposure through lactation^46,47^. In addition, we used 16S amplicon sequencing to characterize the gut bacterial communities. This method is limited in resolution as many recent studies have revealed that host-microbe and microbe-microbe interactions occur at as species and subspecies-level variants^44,45^. Finally, aside from the gut microbiome, various other physiological mechanisms are altered in rodent offspring after exposure to artificial sweeteners *in utero*^21–24^ (i.e. intestinal sugar absorption stimulation, increased postnatal weight gain, altered lipid profiles, downregulation of hepatic detoxification, and increased adulthood insulin resistance). Although we were unable to explore these mechanisms in our study, they will be addressed by future work in the CHILD cohort involving metagenomics of infant stool and metabolomics of infant stool, urine and serum.

## CONCLUSION

In this study, we characterized the infant gut microbiome of 100 infants and found evidence that maternal ASB consumption during pregnancy might have unforeseen effects on infant gut microbiome development and body mass index during the first year of life. As we face an unprecedented rise in childhood obesity and related metabolic diseases, further research is warranted to understand the impact of artificial sweeteners on gut microbiome and weight gain, especially during critical periods of early development.

## ARTICLE INFORMATION

### Author contributions

ABB, PJM, SET, TJM, MRS, and PS, coordinated the CHILD cohort and collected the data; MCA, MBA, and LKS designed the study and obtained funding; ILL and MCA analyzed the data; ILL, MCA, MBA, and LKS interpreted the results, wrote and edited the manuscript. All authors critically reviewed the manuscript and approved the final version for submission.

### Funding/Support

The Canadian Institutes of Health Research (CIHR) and the AllerGen Network of Centres of Excellence provided core support for the CHILD Cohort Study. This research was specifically funded by a CIHR Operating Grant for the Secondary Analysis of Existing Cohort Data (#151530, co-PIs: MBA and MCA).

This research was supported, in part, by the Canada Research Chairs Program. MBA holds a Tier 2 Canada Research Chair in the Developmental Origins of Chronic Disease and is a CIFAR Fellow in the Humans and the Microbiome Program; SET holds a Tier 1 Canada Research Chair in Pediatric Precision Health; ILL hold a Tier 2 Canada Research Chair in Applied Microbial Ecology.

## Acknowledgements

We are grateful to all the families who took part in this study and the whole Canadian Healthy Infant Longitudinal Development Study team, which includes interviewers, nurses, computer and laboratory technicians, clerical workers, research scientists, volunteers, managers, and receptionists. We also thank Alyssa Archibald, MD (University of Manitoba, Winnipeg, Manitoba, Canada), for her assistance with the literature review for this project, and Faisal Atakora, MSc (University of Manitoba), for statistical expertise.

## Data and Code Availability

Raw sequences have been deposited on NCBI public repository (Bioproject #PRJNA624780). The R code, metadata, community matrix and taxa matrix are available on github.

